# CA19-9-associated transcriptional programs in pancreatic ductal adenocarcinoma are linked to survival and drug sensitivity

**DOI:** 10.64898/2026.11.20.767879

**Authors:** Thomas Goralski, Zach Klamer, Chongfeng Gao, Ash Fletcher, Lisa Turner, Tracey Avequin, Katelyn Becker, Chanjuan Shi, Peter Allen, Brian Haab, H. Josh Jang

## Abstract

Pancreatic cancer contains diverse cancer-cell states that complicate treatment selection. We asked whether targeted transcriptomics of glycan-defined PDAC cells would reveal novel and clinically informative subtypes. Spatial profiling of CA19-9-positive regions from 3 tumors with divergent clinical outcomes identified four transcriptional programs: inflammation and tissue remodeling (Cluster 1), proliferation (Cluster 2), antigen presentation and differentiation (Cluster 3), and protein synthesis (Cluster 4). These programs were recovered across 108 Visium sections from 39 patients, 161 patients from The Cancer Genome Atlas, and 37 cell lines with drug-response data. Among 31 tumors with broadly elevated program expression, eight Cluster 1– dominant tumors had poorer survival than the remaining 23. Exploratory drug-response analyses identified cluster-associated sensitivity and resistance patterns, including greater moexipril sensitivity with higher Cluster 1 activity. These findings demonstrate targeted profiling of glycan-defined PDAC populations to uncover cancer-cell-intrinsic therapeutic vulnerabilities, and they reveal CA19-9-associated molecular diversity with program-specific therapeutic candidates.

## Introduction

Pancreatic ductal adenocarcinoma (PDAC) remains among the most lethal malignancies. Surgical resection offers the only potential for cure, yet only 15–20% of patients present with resectable disease, and long-term survival remains limited even after surgery ^1,2^. Large-scale genomic and transcriptomic studies, including The Cancer Genome Atlas, have defined the molecular heterogeneity of PDAC and identified several tumor subtypes^3,4^, although further prospective validation is needed to establish their clinical utility. For example, the predominant PDAC transcriptomic subtypes of classical and basal-like/squamous were associated with differences in survival and responses to FOLFIRINOX and gemcitabine in a phase-II randomized trial^5^.

However, the current subtyping systems have not provided enough predictive accuracy to enable effective precision-medicine strategies for most patients^6^. Transcriptomic classifiers have also shown prognostic value and potential to inform chemotherapy selection^7,8^, although further prospective validation is needed to establish their clinical utility. Recent advances in targeting oncogenic RAS provide proof of principle that cancer-cell-intrinsic dependencies can be therapeutically exploited in PDAC^9^, motivating efforts to identify and target additional cancer-cell intrinsic vulnerabilities specific to distinct tumor subtypes.

Canonical PDAC transcriptomic subtypes, including Classical and Basal-like/Squamous programs, were largely defined through bulk or bulk-deconvolved tumor profiling^10–12^. Although these frameworks have revealed clinically relevant biology, bulk tumor profiles conflate cancer-cell-intrinsic expression with signals from stromal and immune cells^13^. Consequently, therapeutically relevant cancer-cell programs may be detectable in bulk datasets yet remain obscured by sample-level subtype classifications. Single-cell and spatial analyses have demonstrated that classical and basal-like states can coexist within individual tumors, including cells with intermediate or coexpressing phenotypes^14–16^.One such study identified 7 distinct lineage programs that further subdivide among 12 different cell-state programs^17^. Other expression studies have identified multiple cancer cells across the epithelial-mesenchymal spectrum that potentially provide predictive value relating to metastatic capability and treatment resistance^18,19^. Spatially resolved profiling can address this limitation by defining molecular programs directly within phenotypically identified cancer-cell populations^20,21^. While these approaches have revealed the rich complexity of tumor biology within the cancer-cell population and among the non-cancer components, the use of such expression signatures as the basis for prediction of tumor treatment responses has yet to be established. Once defined, these cancer-cell-specific programs can be projected into larger bulk transcriptomic datasets to determine whether they remain detectable despite tissue admixture and whether their relative representation is associated with clinical outcomes and therapeutic vulnerabilities.

Cell-surface glycan patterns, including the clinically established biomarker CA19-9, provide a means of identifying and molecularly characterizing distinct PDAC cancer-cell populations.The cancer-cell glycome is shaped by coordinated changes in transcription^22^, metabolism^23^, and biosynthetic enzyme activity and substrate availability^24^, thereby providing a readout of the cellular state. Glycan alterations are particularly prominent in pancreatic cancer, reflecting both the extensive glycosylation characteristic of pancreatic tissue and the sensitivity of glycosylation to tumor-associated metabolic and transcriptional changes^25^. Integrated glycomic and transcriptomic studies have linked altered glycosylation machinery to pancreatic tumor development, while analyses of human PDAC have identified glycan-associated differences among transcriptional states and their interactions with immune cells^26–28^. The established cancer biomarker CA19-9 is a glycan motif present on mucins and glycosphingolipids that is shed into the blood circulation upon tissue disruption^29^. Additional glycan alterations in PDAC include ABO glycans ^30^, Lewis blood-group structures such as sialy-Lewis X^30,31^, and alpha 2,6-linked sialic acid^32,33^. A glycan that is related to CA19-9, termed sTRA, defines a separate subpopulation of cancer cells^34^ and forms a blood biomarker that complements CA19-9 to provide added diagnostic value^35^. These sTRA-type and CA19-9-type tumors previously were associated with differing responses to chemotherapy and associations with outcomes^35^.

In previous work, we demonstrate that signatures of 4 glycan biomarkers distinguished PDAC cancer-cell populations and stratified primary tumors into three types: sTRA-type, CA19-9-type, and intermixed ^36^. These glycan-defined populations provide a clinically relevant basis for selecting and molecularly interrogating specific cancer-cell populations within intact PDAC tumors. Additional work indicated functional differences between the groups, as the CA19-9-type cells were enriched for epithelial and classical features, whereas the sTRA-type cells encompassed both epithelial/classical and mesenchymal/basal-like phenotypes showed higher chemotherapy resistance and associations with shorter survival^37^. Among these glycan-defined populations, CA19-9-positive cells are particularly relevant because CA19-9 remains the most widely used PDAC-associated biomarker. Beyond its biomarker role, CA19-9 has been shown to promote pancreatic inflammation and tumorigenesis in mouse models^38^, its role in metastasis as a ligand for E-selectin^39^, and its potential value as an immunotherapy target ^40^. These findings suggest that the glycan-defined populations provide a clinically relevant basis for guiding targeted transcriptional studies within heterogeneous PDAC tumors.

Therefore, we asked whether targeted transcriptomics of glycan-defined PDAC cell subpopulations reveals informative transcriptional subtypes that would otherwise be missed by bulk studies or untargeted discovery methods. We hypothesized that CA19-9-positive cells harbor distinct transcriptional programs that can be recovered across independent datasets and linked to clinically relevant phenotypes. We used the novel approach of guiding the selections of cancer cells through multiplexed glycan immunofluorescence detection, enabling spatial transcriptomic and proteomic profiling of CA19-9-positive PDAC cells from patients with divergent clinical outcomes. We used GeoMx profiling on three patient PDAC tumors to define expression programs within marker-selected CA19-9-positive cancer-cell compartments. We then projected these programs into an independent PDAC Visium spatial transcriptomic cohort and the TCGA-PAAD bulk RNA-sequencing cohort and evaluated their associations with drug response in PDAC cell lines. By separating malignant-cell programs from surrounding tissue signals, this approach revealed cancer-cell-intrinsic transcriptional heterogeneity within CA19-9-positive PDAC. These spatially defined programs could be recovered in larger datasets for associations with patient outcomes and candidate therapeutic vulnerabilities. Together, our findings support the value of guided spatial profiling of glycan-defined PDAC populations to uncover cancer-cell-intrinsic heterogeneity and therapeutic vulnerabilities that would be obscured by bulk tissue or untargeted analyses.

## Results

### CA19-9–positive PDAC samples share a glycan-defined state but differ in transcriptomic program

Our previous work defined PDAC tumors as being derived from distinct glycan biomarker defined lineages, with some tumors having a clear CA19-9 only population, others having a sTRA only population, and the remaining belonging to a mixed lineage.^36^We selected adjacent sections from the same tissue blocks analyzed in our previous glycan-profiling study, selecting three tumors that all belonged to the CA19-9-defined lineage. Despite this shared glycan identity and broadly similar clinical features, the patients had divergent clinical outcomes: two developed early recurrence, whereas the third has remained without evidence of disease for more than six years. This survival disparity among patients within a shared CA19-9–defined tumor class motivated us to ask whether CA19-9–positive PDAC contains distinct cancer-cell transcriptional programs that reflect biological heterogeneity and variable clinical response, with the goal of identifying molecular vulnerabilities that can be further personalized for patients with these glycan-decorated PDAC tumors.

To determine whether CA19-9–positive PDAC harbors a uniform cancer-cell transcriptional program or multiple distinct oncogenic programs, we used GeoMx Digital Spatial Profiling platform to analyze regions of interest drawn around tumor crypts composed predominantly of ductal epithelial cancer cells (**Fig. 1**). Selected regions were profiled for both whole-transcriptome RNA expression using the GeoMx Whole Transcriptome Atlas and protein expression using the GeoMx IO Proteome Atlas. Cancer cell purity and glycan expression in these tumor regions were independently validated by manual multiplex immunofluorescence and expert pathologic review. This strategy allowed us to directly link a glycan-defined tumor states with spatially resolved cancer-cell molecular programs.

**Figure 1.**
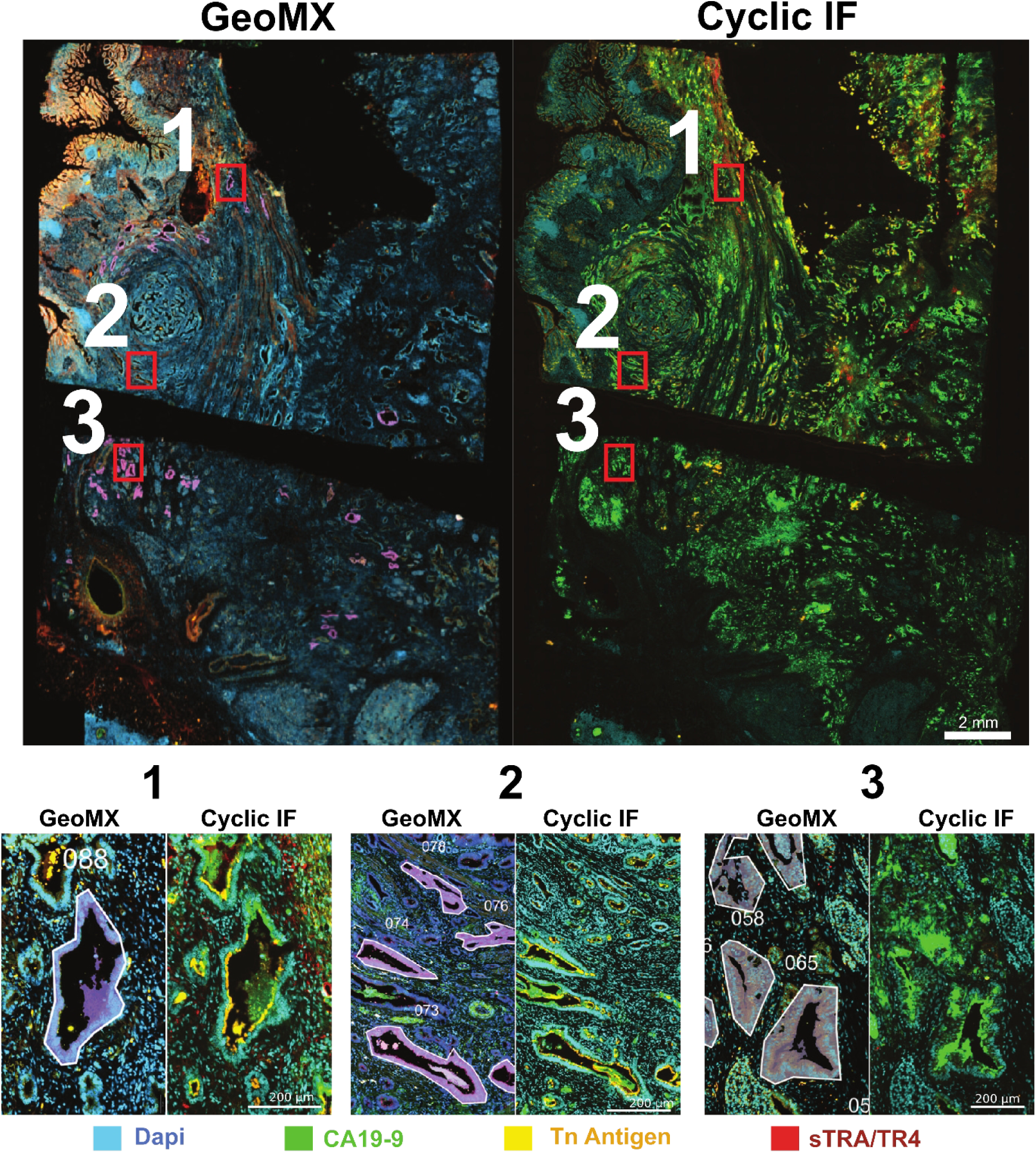
Glycan-directed selection of pancreatic ductal adenocarcinoma regions for GeoMx spatial profiling. Representative cyclic immunofluorescence images of a PDAC tissue section stained for nuclei (DAPI, cyan), CA19-9 (green), Tn antigen (yellow), and sTRA/TRA (red). The GeoMX segmentation overlay (White) overlays these colors. Whole-section views show the distribution of glycan-defined tumor populations, with numbered image boxes identifying three representative regions enlarged below. In the corresponding higher-magnification images, outlined areas indicate cancer-enriched regions of interest and areas of illumination selected for GeoMx whole-transcriptome and protein profiling. Scale bars are shown in the images.

A total of 172 areas of illumination (AOIs) from three tissue specimens were profiled. After quality control, 164 AOIs (95.3%) were retained for matched RNA and protein analyses, comprising 75 AOIs from BRPC_17-543, 43 from BRPC_19-296, and 46 from BRPC_20-281. PCA dimension reduction of cancer-cell RNA expression revealed separation among the three CA19-9–positive samples, indicating that tumors sharing CA19-9 positivity nevertheless harbored distinct transcriptomic identities (**Fig. 2B)**. After regression of between-subject variation, the dominant sample-level separation was reduced, supporting the interpretation that either patient identity or tumor-to-tumor heterogeneity contributed to the expression differences (**Fig. 2B, Supp. Fig. 1A**).

**Figure 2.**
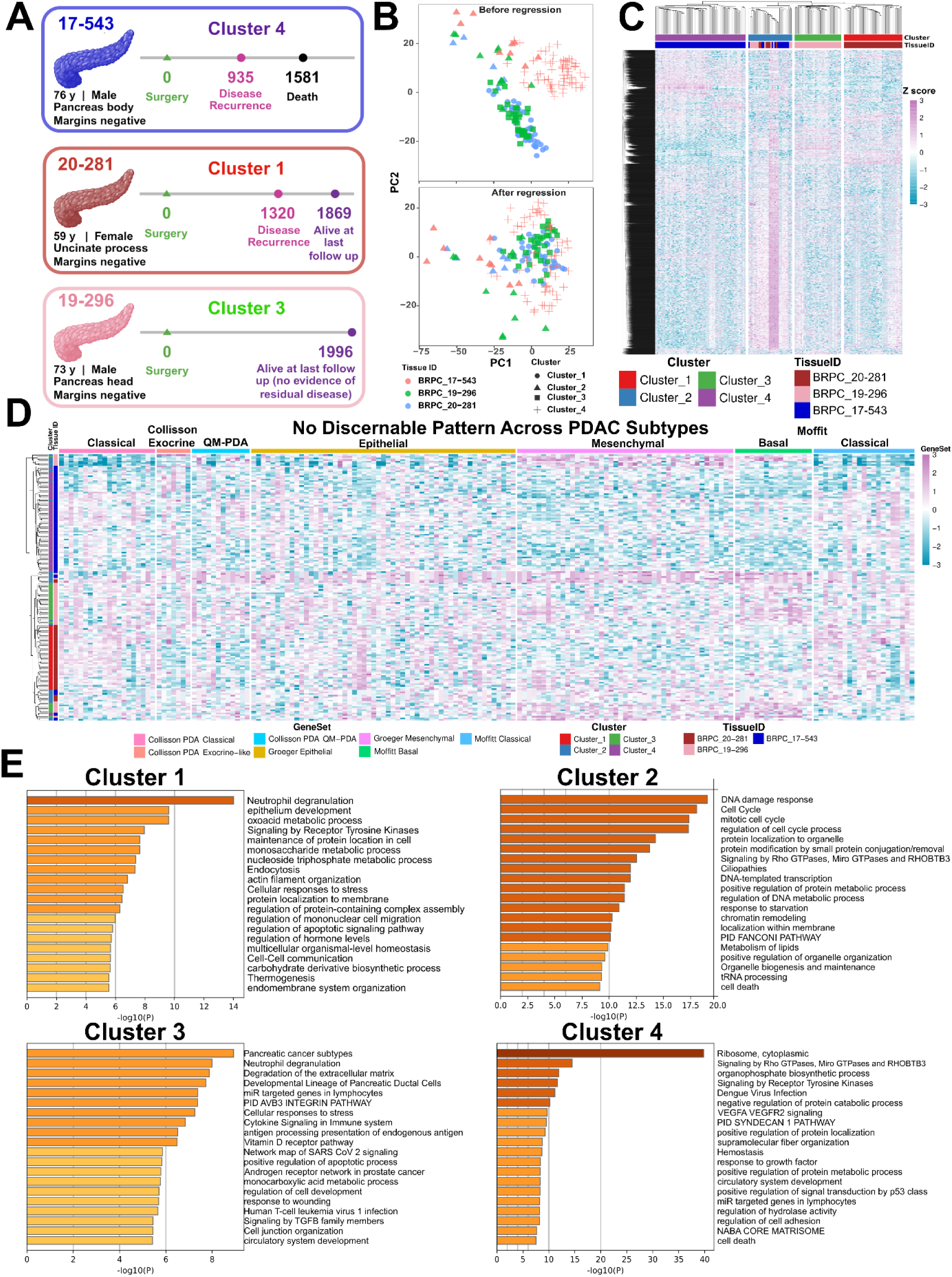
Discovery of CA19-9-associated transcriptional programs in PDAC. **(A)** Clinical overview of the three CA19-9–positive PDAC cases profiled by GeoMx, showing the predominant expression program associated with each tumor and the timing of surgery, recurrence, and death or last follow-up. **(B)** Principal component analysis of normalized RNA expression across GeoMx areas of illumination (AOIs), colored by patient and shaped by expression cluster. The upper panel shows unadjusted expression data, whereas the lower panel shows expression after regression of patient-level effects. **(C)** Unsupervised hierarchical clustering of global GeoMx RNA expression identified four transcriptional programs across the profiled AOIs. Columns represent AOIs and rows represent genes; annotations indicate patient identity and RNA-cluster assignment. **(D)** Characterization of the four GeoMx-derived programs using published Classical and Basal-like/Squamous PDAC signatures, glycosylation-associated genes, and Gene Ontology enrichment. Enriched biological processes included inflammatory and tissue-remodeling programs in Cluster 1, DNA-replication and cell-cycle programs in Cluster 2, antigen-presentation, differentiation, and stress-response programs in Cluster 3, and ribosomal assembly and translation programs in Cluster 4.

Unsupervised clustering of global RNA expression further resolved multiple expression programs across the GeoMx regions (**Fig. 2C**). These programs were associated with sample identity but showed structured, reproducible gene-expression patterns rather than random variation. Therefore, we first asked if these reproducible patterns reflected previously established PDAC transcriptional subtypes. Importantly, the observed organization was not explained by canonical Classical and Basal-like/Squamous PDAC subtype features, suggesting that CA19-9-associated transcriptional heterogeneity represents an additional layer of tumor-state variation between or across established subtype frameworks (**Fig. 2D**).

To define the biological processes underlying this subtype-independent heterogeneity, we next performed Gene Ontology analysis of cluster-defining genes, which revealed distinct functional enrichments among the 50 highest-ranking terms for each cluster (**Supp. Fig. 2**). Cluster 1 was enriched for leukocyte and macrophage migration, wound healing, and actin remodeling programs, suggesting an inflammatory and tissue-remodeling-associated cancer-cell state or potentially a more invasive/metastatic phenotype. Cluster 2 was enriched for DNA biosynthesis, G2/M cell-cycle, nuclear division, and chromatid segregation pathways, consistent with a proliferative program shared across CA19-9-positive tumors. Cluster 3 was enriched for antigen processing and MHC-related pathways, endodermal differentiation, TGF-β signaling, and apoptotic/endoprotease regulation, suggesting a program linked to differentiation, immune presentation, and stress-response biology.

Cluster 4 was dominated by ribosomal small-subunit assembly and cytoplasmic translation programs, indicating a translational or biosynthetic emphasis. These enrichments support the interpretation that the RNA-defined clusters reflect biologically distinct transcriptional programs within the CA19-9-positive lineage. The patient with tumor associated with Cluster 4 succumbed to disease in 1581 days, patient with tumor associated with Cluster 1 had disease recurrence at day 1320, and patient with tumor associated with Cluster 3 had progression free survival tracked through 1996 days (**Fig. 2A**).

Because CA19-9 is a glycan-defined phenotype, we next asked whether the RNA-defined programs were accompanied by variation in glycosylation-related transcriptional pathways. Glycan-related genes showed coordinated differences across the same sample/program architecture, linking the CA19-9-positive state to differences in the transcriptional regulation of glycan biology (**Supp. Fig. 1C**). Protein-level GeoMx profiling similarly revealed structured differences across samples and expression programs, providing an orthogonal readout that the RNA-defined profiles correspond to broader phenotypic differences (**Supp. Fig. 1B**).

Together, these data show that CA19-9 positivity does not define a single uniform cancer-cell transcriptomic state. Instead, CA19-9–positive PDAC samples exhibit distinct cancer-cell programs, including inflammatory/remodeling, proliferative, immune/differentiation-associated, and translational states, accompanied by variation in glycan-related gene and protein expression.. These patient-derived CA19-9-associated profiles therefore provided a discovery framework for testing whether spatially defined cancer-cell programs can be recovered across independent PDAC datasets and leveraged to reveal clinically relevant biology and candidate therapeutic vulnerabilities.

### GeoMx-derived CA19-9-associated programs detected in independent PDAC Visium cohort

To test whether the GeoMx-derived CA19-9-associated programs could be detected in an orthogonal spatial transcriptomic platform, we projected the transcriptional cluster signatures onto an independent Visium PDAC cohort^41^. In this dataset, transcriptomes from 39 PDAC tumors and 108 sections were available, resulting in 353,257 spots in total. 154,169 cancer-enriched Visium spots were identified from spatial annotations, allowing comparison of GeoMx-derived cancer-cell programs across a spot-based transcriptomic dataset with different spatial resolution and tissue-processing features (**Fig. 3A**).

**Figure 3.**
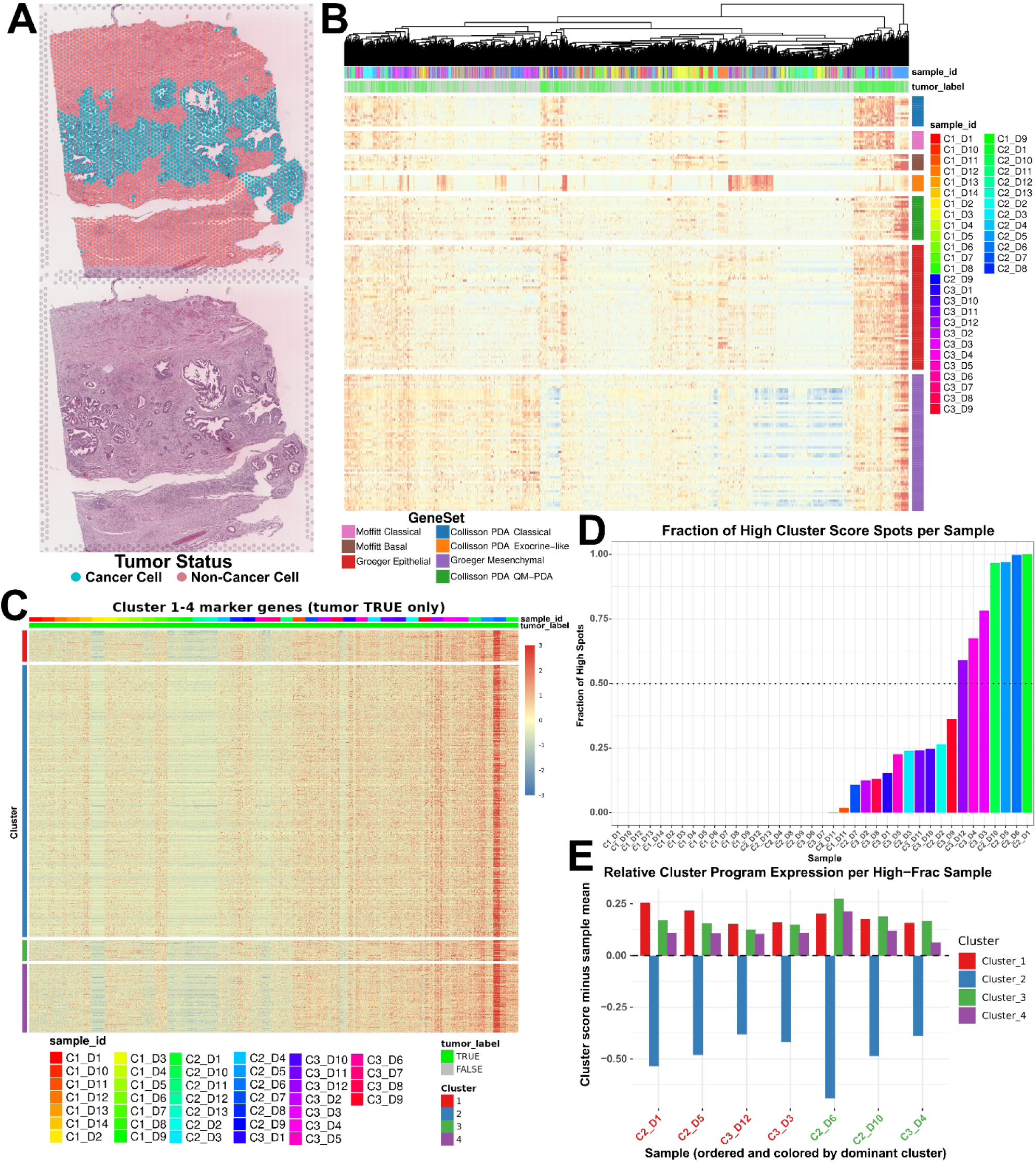
Recovery of GeoMx-derived CA19-9-associated transcriptional programs in an independent PDAC Visium cohort. **(A)** Representative tissue section showing inferred cancer-enriched spots overlaid on the hematoxylin-and-eosin image (top) and the corresponding image without spot annotations (bottom). Cancer enrichment was inferred from a weighted combination of cancer, noncancer, and immune module scores, averaged over the six nearest spatial neighbors. Spots with smoothed scores above the section mean were classified as cancer-enriched. **(B)** Expression of established PDAC subtype and epithelial/mesenchymal marker genes across randomly sampled Visium spots (up to 200 per sample). Rows represent spots ordered by hierarchical clustering; columns represent genes grouped by published gene set. Annotations indicate sample identity, tumor classification, and gene-set membership. Colors represent gene-wise standardized expression. **(C)** Expression of marker genes from the four GeoMx-derived CA19-9-associated clusters across tumor-labeled Visium spots (up to 150 randomly selected spots per sample). Rows represent genes grouped by their originating GeoMx cluster; columns represent spots grouped by sample, with samples ordered by the fraction of high-scoring spots shown in panel D. Colors represent gene-wise standardized expression, and annotations indicate cluster membership, sample identity, and tumor classification. **(D)** Fraction of high-scoring spots in each Visium sample, ordered from lowest to highest. Each spot was scored by calculating the mean normalized expression of available marker genes within each GeoMx cluster and then averaging the four cluster scores. Spots with an average score greater than 2.214 were designated high-scoring. The dotted line marks a fraction of 0.5. **(E)** Relative cluster scores in samples with a high fraction (High-Frac) of high-scoring spots. Bars show each sample’s cluster scores after subtraction of the sample mean across the four clusters, highlighting relative program expression. Samples are ordered by their highest-scoring cluster, and sample labels are colored by that dominant cluster.

Spot-level expression analysis revealed broad transcriptional heterogeneity across cancer-enriched regions in the PDAC Visium cohort. Consistent with the GeoMx discovery analysis, canonical PDAC subtype signatures did not clearly segregate cancer-enriched spots. Instead, pathway-associated programs varied across samples and spatial regions, revealing diverse molecular states not captured by bulk subtype classifications (**Fig. 3B**). Pathway-level scoring, including analysis of glycolysis-associated programs, further demonstrated spatially organized functional heterogeneity among cancer-enriched regions within individual tumors (**Supp. Fig. 3C**).These features provided a stringent context in which to evaluate whether the GeoMx-derived CA19-9-associated programs could be recovered outside the original discovery samples.

Projection of the four GeoMx-derived programs onto cancer-enriched Visium spots revealed strong correlations among program scores (**Supp. Fig. 3A)**, indicating that the signatures capture a substantial shared transcriptional component in this dataset. Because CA19-9 expression was not directly measured in these tumors, this common component was interpreted as a GeoMx-profile-like or tumor-epithelial signal rather than definitive evidence of CA19-9 positivity. Also, the strong correlation among absolute cluster program scores limited their ability to distinguish cluster program-specific activity. We therefore used the mean score across all four programs to quantify overall recovery of the GeoMx-derived profile and relative program scores to assess the dominant program emphasis within profile-high spots and samples (**Fig. 3D**).

The distribution of mean cluster scores showed that GeoMx-profile-like expression was not uniformly present across the Visium PDAC samples. Instead, most spots showed low-to-intermediate scores, while a subset of spots showed high similarity to the GeoMx-derived programs (**Fig. 3C, Supp Fig. A-B**). At the sample level, only a subset of tumors contained a high fraction (>50%) of profile-high spots, whereas many samples showed little evidence of recovery by the current discovery signatures (**Fig. 3C-D**). Among high-identity Visium samples, GeoMx-profile-like genes retained structured expression patterns, supporting recovery of the discovery marker architecture rather than stochastic expression of individual genes (**Fig. 3C**). Relative cluster scoring further revealed distinct dominant program emphases among high-fraction samples, with some tumors showing stronger Cluster_1-like activity and others showing stronger Cluster_3-like activity (**Fig. 3E**). Together, these data demonstrate cross-platform detection of CA19-9-associated transcriptional programs in an orthogonalVisium PDAC cohort while also emphasizing the need for expanded glycan-linked spatial profiling and clinical outcome annotation to determine how broadly these programs generalize across PDAC.

### Dominant CA19-9-associated programs in TCGA-PAAD are linked to patient survival

Having identified distinct CA19-9-associated cancer-cell expression profiles in GeoMx and Visium PDAC data, we next asked whether these transcriptional signatures were detectable in bulk PDAC tumor transcriptomes and whether this was associated with clinical outcome. We projected the GeoMx-derived cluster signatures onto TCGA-PAAD bulk tumor RNA-seq data. In total, there were 161 PDAC patient tumors included in the bulk transcriptomes. For each TCGA tumor sample, the four cluster scores were calculated as the mean FPKM-UQ expression of the available marker genes assigned to each GeoMx-derived cluster. Tumors were classified as CA19-9-like if both the mean and minimum of their four cluster scores met or exceeded the respective 75th-percentile thresholds across the analyzed TCGA-PAAD samples.This criterion selected tumors with broadly elevated expression across the four CA19-9-associated programs, rather than tumors with high expression confined to a single program.This generated four cluster identity scores per TCGA tumor, corresponding to the four CA19-9-associated transcriptomic signatures identified in the GeoMx discovery cohort. Despite the loss of spatial resolution and the presence of stromal and immune admixture in bulk tissue profiles, these scores showed structured variation across TCGA tumors, indicating that transcriptional features associated with CA19-9-glycan-positive PDAC regions remain detectable at the cohort level, but may have been obscured or underappreciated in conventional bulk analyses. (**Fig. 4A**).

**Figure 4.**
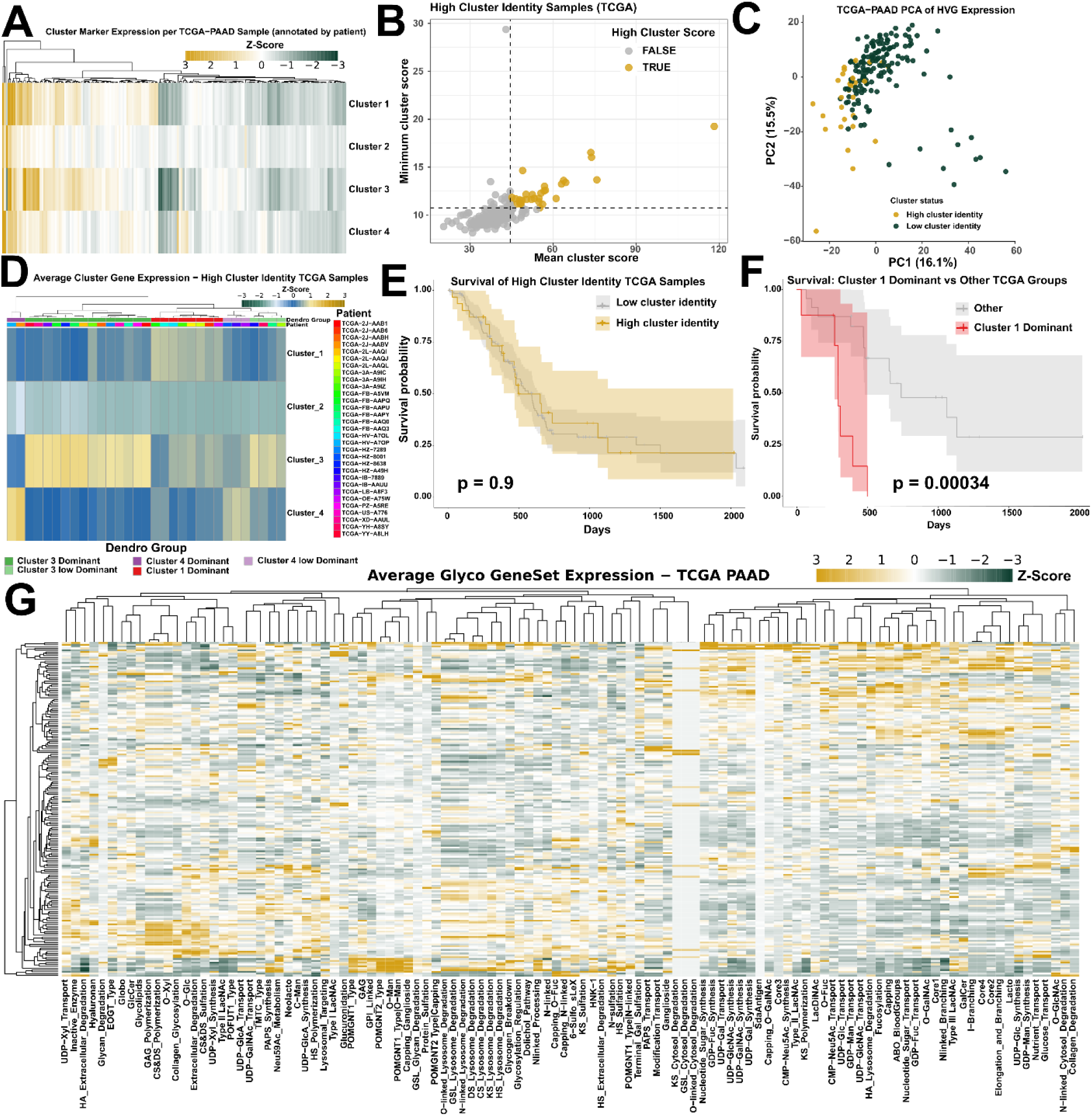
Projection of GeoMx-derived CA19-9-associated programs into TCGA-PAAD identifies a poor-outcome Cluster 1–dominant tumor group. **(A)** Heatmap of the four GeoMx-derived program scores across TCGA-PAAD tumors. For each tumor, each program score was calculated as the mean upper-quartile-normalized FPKM expression of the available marker genes assigned to that program. Columns represent TCGA samples and rows represent GeoMx programs; samples were ordered by hierarchical clustering. **(B)** Classification of TCGA tumors by overall GeoMx-profile identity. Each point represents one tumor, positioned according to its mean score across the four programs and its minimum individual program score. Dashed lines indicate the respective 75th-percentile thresholds. Tumors meeting or exceeding both thresholds were classified as high-profile-identity tumors and are shown in gold. **(C)** Principal component analysis of TCGA-PAAD transcriptomes using the 1,000 most variable genes. Points represent tumors and are colored according to high- or low-profile-identity status. **(D)** Within-sample-scaled expression of the four GeoMx-derived programs among high-profile-identity tumors. Columns represent tumors and rows represent programs. Hierarchical clustering of relative program expression was used to identify groups with distinct dominant program emphasis; the upper annotation indicates the assigned dominant-program group. **(E)** Kaplan–Meier overall-survival analysis comparing high-profile-identity tumors with all other TCGA-PAAD tumors. Survival time was defined as days to death or last follow-up, and groups were compared using a two-sided log-rank test. Shaded regions indicate 95% confidence intervals. **(F)** Kaplan–Meier overall-survival analysis comparing Cluster_1–dominant tumors with the other dominant-program groups among high-profile-identity tumors. Cluster_1 dominance was associated with shorter overall survival by two-sided log-rank testing. Shaded regions indicate 95% confidence intervals. **(G)** Heatmap of glycosylation-associated gene-set scores across TCGA-PAAD tumors. Scores were calculated as the mean expression of genes available within each gene set and scaled across tumors by gene set. Rows represent glycosylation-related gene sets and columns represent tumors.

To identify TCGA tumors with coordinated representation of the GeoMx-derived signatures, we summarized each tumor using the mean of its four program scores and the minimum score across the four programs (**Fig. 4B**). Samples with both high mean score and high minimum score were classified as high cluster-identity tumors, totaling 31 high cluster identity tumors out of 161 total tumors. This dual-threshold strategy selected tumors with coordinated expression across all four GeoMx-derived programs, rather than isolated elevation of a single program, thereby enriching for tumors that broadly recapitulated the transcriptional features of CA19-9-positive regions.

Principal component analysis further showed that high-profile-identity tumors occupied a partially distinct region of global transcriptomic space, supporting their classification as a transcriptionally coherent subset within TCGA-PAAD (**Fig. 4C**). To determine whether this structure was specific to the GeoMx-derived marker genes and the high cluster-identity subset, rather than an artifact of small-sample selection, we performed 1,000 simulation-based resampling iterations. In each iteration, three TCGA tumors were sampled from either the high- or low-identity pool and scored using either the authentic GeoMx marker sets or size-matched random gene sets. High cluster-identity tumors scored with the authentic markers showed stronger recovery of the reference program structure than the comparison conditions, supporting the nonrandom transferability of the GeoMx-derived signatures (all adjusted *P*<0.0001; **Supp. Fig. 4B**).

Survival analyses included 161 patients with available survival information, comprising 31 tumors classified as CA19-9-like and 130 classified as profile-low. Among the 31 CA19-9-like tumors, 8 were Cluster 1–dominant and were compared with the remaining 23 tumors to evaluate the association between Cluster 1 dominance and survival. High cluster-identity status alone was not significantly associated with patient survival probability (**Fig. 4E**), indicating that broad representation of the CA19-9-associated profiles is not, by itself, sufficient to define a uniformly aggressive or protective disease state. However, high cluster-identity tumors were not transcriptionally uniform, and bulk TCGA expression profiles are influenced by non-malignant stromal and immune transcriptional signatures. We therefore examined the relative representation of the four GeoMx-derived programs within each high-profile-identity tumor. Program scores were scaled within each tumor and hierarchically clustered to identify groups with distinct dominant program emphases (**Fig. 4D**). This classification reflected which program was preferentially represented relative to the other programs within each tumor. We next asked whether dominant program identity, rather than high-profile-identity status alone, was associated with patient survival.. Kaplan–Meier analysis showed that tumors with Cluster_1-dominant profile identity, which represented inflammatory and tissue-remodeling signatures, had significantly worse survival than other TCGA-PAAD tumors (**Fig. 4F, Supp. Fig. 4A**). Thus, the prognostic association emerged only after stratifying high-profile-identity tumors by dominant program identity, demonstrating that tumors with similar overall representation of the GeoMx-derived CA19-9-associated profiles differed in survival according to their relative transcriptional program composition.Finally, glycan-related gene-set expression showed structured variation across TCGA-PAAD tumors, linking the recovered GeoMx-derived profile structure to broader glycosylation-associated transcriptional biology (**Fig. 4G**). Together, these results show that transcriptional features distinguishing CA19-9-glycan-positive GeoMx profiles are detectably structured in TCGA bulk RNA-seq and that Cluster 1-dominant profile identity identifies a poor-prognosis subset of TCGA-PAAD tumors.

### CA19-9-associated transcriptional programs stratify drug-response phenotypes in PDAC cell lines

Having discovered that specific CA19-9-associated transcriptional signatures were associated with patient survival in TCGA-PAAD, we next asked whether these programs were linked to pharmacologic response in experimentally tractable *in vitro* PDAC models. We projected the GeoMx-derived cluster signatures onto DepMap-profiled pancreatic cancer cell lines with matched RNA-seq and drug-sensitivity data. Of 59 pancreatic cancer cell line models identified in DepMap, 37 had both transcriptomic and pharmacologic response data acros 4517 distinct drugs, and 4685 distinct drug-dose combinations which were retained for integrated analysis.

We performed GeoMx cluster scoring, which revealed that the CA19-9-associated programs were differentially represented across DepMap pancreatic cancer cell lines (**Fig. 5A**). Cluster_2, which was the most consistently shared program across patient tumors, did not define a distinct high-scoring cell-line subgroup after relative scoring, consistent with its interpretation as a common CA19-9-associated cell cycle component rather than a cancer-subtype program. In contrast, Cluster_1, Cluster_3, and Cluster_4 showed greater cell-line-to-cell-line variation, suggesting that these programs capture more specific cell-state transcriptional signatures in the DepMap panel. This provided a framework to test whether continuous program activity, rather than categorical subtype assignment alone, was associated with drug-response phenotypes.

**Figure 5.**
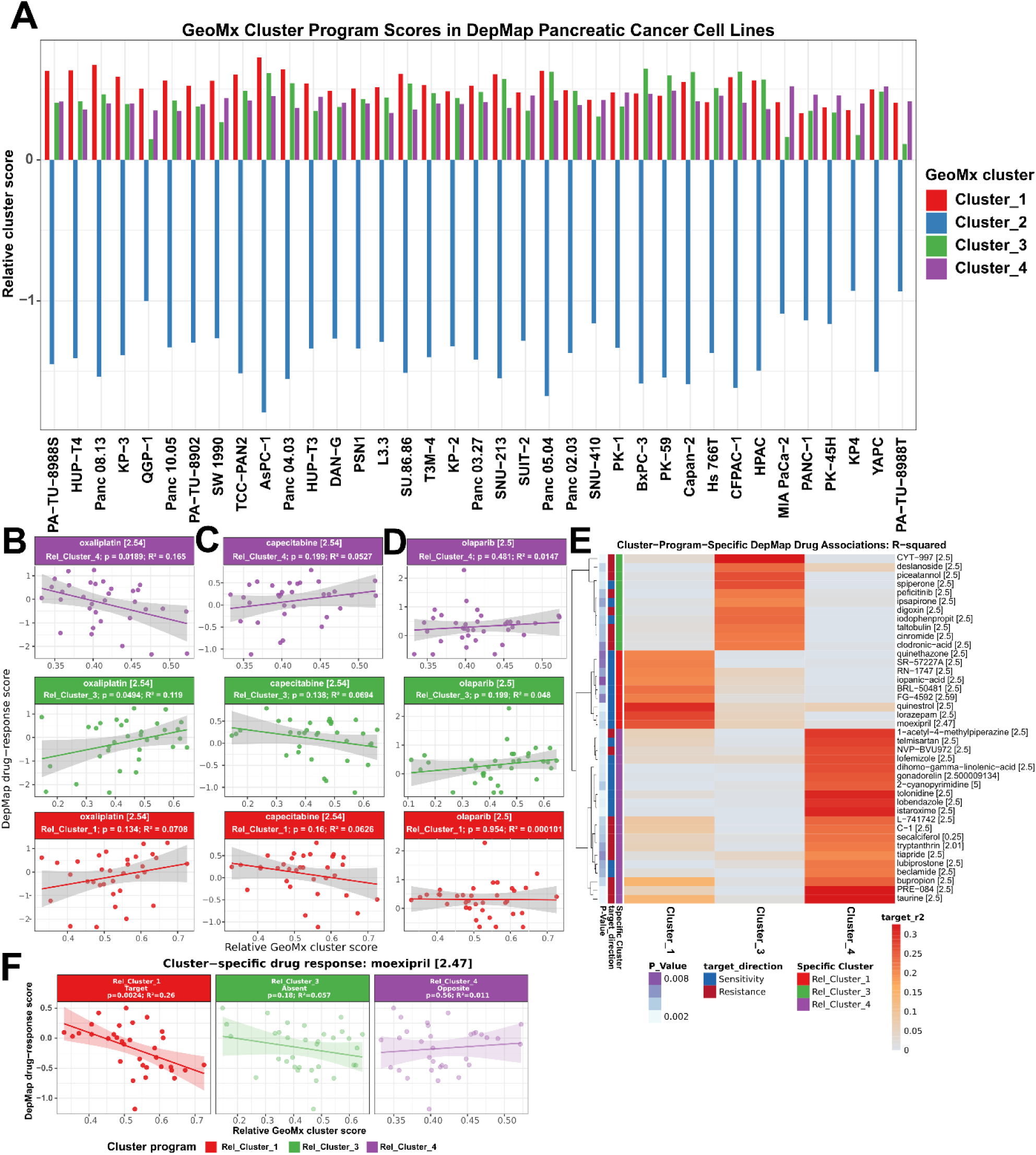
GeoMx-derived CA19-9-associated programs identify drug-response phenotypes in pancreatic cancer cell lines. **(A)** Relative GeoMx program scores across 37 DepMap pancreatic cancer cell lines with matched RNA-sequencing and pharmacologic-response data. For each cell line, the mean score across the four programs was subtracted from each individual program score; positive values indicate relative enrichment and negative values indicate relative depletion within that cell line. Cell lines are ordered by their dominant program. Cluster 2 showed comparatively uniform representation and was therefore not emphasized in subsequent program-specific analyses. **(B–D)** Associations between relative GeoMx program scores and DepMap response to selected standard-of-care or mechanistically related agents. Each point represents a pancreatic cancer cell line. Lines and shaded regions show ordinary least-squares regression estimates and 95% confidence intervals, respectively; drug, dose, nominal *P* value, and R² are displayed within each facet. More negative DepMap response values indicate greater sensitivity; therefore, negative slopes indicate increasing sensitivity with increasing program score, whereas positive slopes indicate relative resistance. **(E)** Heatmap of prioritized cluster-program-specific drug associations from the broader DepMap pharmacologic screen. Rows represent drug–dose pairs and columns represent relative Cluster 1, Cluster 3, and Cluster 4 scores. Color intensity represents the R² from the corresponding continuous regression model. Candidate associations required a nominal target-program *P*<0.05, target-program R²≥0.20, and no conflicting same-direction association of comparable magnitude in the other programs. Row annotations indicate the program showing the strongest association and whether the association predicted sensitivity or resistance. **(F)** Moexipril response modeled against relative Cluster 1, Cluster 3, and Cluster 4 program scores. Higher Cluster 1 scores were associated with more negative response values, consistent with greater moexipril sensitivity, whereas the other programs showed weaker or discordant associations. Points represent cell lines; lines and shaded regions show ordinary least-squares regression estimates and 95% confidence intervals. These pharmacogenomic analyses are exploratory and are presented with effect sizes, R² values, and nominal and multiplicity-adjusted statistical results as indicated.

Because the GeoMx-derived programs varied continuously across cell lines, we modeled drug response using regression rather than discrete subtype labels, preserving graded variation in program activity. Power analysis indicated that the matched cohort of 37 cell lines provided at least 80% power to detect associations explaining 20% of drug-response variance (R² = 0.20) at a nominal α of 0.05 (**Supp. Fig. 6**). After Bonferroni correction across the modeled associations, however, achieving 80% power required 134 cell lines for R² = 0.20 and 42 cell lines even for R² = 0.50. Thus, although the cohort could identify associations at nominal significance, it had less than 80% power across the full range of effect sizes evaluated (R² = 0.01–0.50) after correction. We therefore interpreted individual associations as exploratory and emphasized effect size, directionality, and program specificity alongside nominal and multiplicity-adjusted statistical results.

We modeled DepMap drug-response scores as a function of relative GeoMx cluster program scores. Because more negative DepMap drug-response scores indicate greater drug sensitivity, negative regression slopes indicate that higher program scores are associated with increased drug sensitivity, whereas positive slopes indicate reduced sensitivity or increased resistance. Across standard-of-care and related drugs or therapeutic agents, Cluster_1 scores did not show a uniform association with enhanced sensitivity (**Fig. 5B–E, Supp. Fig. 5**).

In contrast, Cluster 3 and Cluster 4 program scores showed more drug-informative patterns, with several associations suggesting selective sensitivity or resistance to specific agents, including standard-of-care therapeutic compounds (E.g. 5-Fluorouracil, SN-38, capecitabine, cisplatin, gemcitabine, inrinotecan, leucovorin, olaparib, oxaliplatin, and paclitacel). For example, Cluster 4-associated trends were observed for irinotecan-related response, where a negative association would indicate greater sensitivity among cell lines with higher Cluster 4 program scores. Next, to identify candidate therapies with stronger subtype-specific relationships than the standard-of-care agents, we next used the broader DepMap drug-response dataset to prioritize certain drug targets for CA19-9-related PDAC tumors. For each GeoMx-derived program, we selected drugs with the largest program-specific effect sizes, allowing us to nominate compounds whose response profiles were preferentially associated with Cluster_1, Cluster_3, or Cluster_4 activity (**Fig. 5E**). Therefore, we identify high-effect candidate vulnerabilities within each CA19-9-associated transcriptional program that can be further explored and evaluated in future studies. In particular, Moexipril was a clinically promising example from this prioritization strategy. For this drug, a higher Cluster_1 program score was associated with more negative DepMap response values, consistent with increased sensitivity among Cluster_1-like PDAC cell lines (**Fig. 5F**). Together, these findings suggest that certain CA19-9-associated programs may encode weak but detectable response differences to standard-of-care treatments, whereas Cluster 1-like tumors may require alternative therapeutic strategies beyond standard cytotoxic agents.

## Discussion

In this study, we leveraged spatially resolved molecular profiling to identify and evaluate cancer-cell-intrinsic expression programs in PDAC that may be obscured by stromal and immune admixture in bulk tissue analyses. We further examined whether PDAC cells sharing a glycan-defined phenotype harbor distinct transcriptional states associated with patient survival and differential drug response. Even with limited patient samples, we were able to identify transcriptomic heterogeneity within CA19-9–positive PDAC and show that this glycan-defined tumor population is not a single uniform transcriptomic state. Spatial profiling of CA19-9– positive tumor regions identified distinct cancer-cell expression programs, with corresponding differences in glycan-related transcriptional features and protein expression. Projection of these programs into independent datasets demonstrated that the GeoMx-derived profiles are recoverable across bulk RNA-seq, cell-line transcriptomes, and Visium spatial transcriptomics. Importantly, clinically and functionally relevant phenotypes were linked to the dominant transcriptional program rather than overall CA19-9-associated profile identity.

These findings extend current models of PDAC heterogeneity. Previous studies have shown that PDAC transcriptional states form a continuum and can coexist within individual tumors, with their representation influenced by both genomic alterations and the microenvironment^14–16^. Canonical Classical and Basal-like/Squamous subtypes have provided a foundational framework for understanding PDAC biology, but these categories were largely derived from bulk or bulk-deconvolved transcriptomic profiles. As a result, they can compress spatially restricted malignant-cell states, mixed tumor composition, and biomarker-defined phenotypes into sample-level classifications. Previous glycomic and transcriptomic studies have linked glycan features to PDAC transcriptional states and immune interactions^27,28^. Our findings extend this work by resolving distinct transcriptional programs within a shared CA19-9-positive phenotype, supporting glycan-defined identity as a complementary layer of tumor stratification.. Thus, CA19-9 positivity should not be interpreted as a uniform malignant-cell phenotype, but rather as a clinically accessible glycan feature that can be associated with multiple cancer-cell transcriptional states. Using an orthogonal spatial transcriptomic platform, we recovered GeoMx-derived program signatures in an independent PDAC Visium cohort. Canonical bulk-defined PDAC subtype signatures did not clearly organize cancer-enriched regions; instead, the Visium data revealed substantial spatial transcriptional heterogeneity within and across tumors.These observations are consistent with spatial studies linking malignant-cell identity to local tissue organization and microenvironmental composition^20,21,42^.

The TCGA analysis highlights the importance of separating overall CA19-9-associated profile identity from dominant program identity. Tumors with high overall similarity to the GeoMx-derived CA19-9-associated profiles did not show a significant survival difference compared with other TCGA-PAAD tumors. However, among tumors with strong profile identity, dominant program emphasis was clinically informative, with Cluster_1 dominance associated with worse survival. This distinction argues against a simple model in which CA19-9-associated identity itself is uniformly aggressive. Instead, prognosis appears to depend on the specific transcriptional state present within the broader CA19-9-associated phenotype. The inflammatory and tissue-remodeling features of Cluster 1 also raise questions about how glycan-associated cancer-cell states interact with the immune microenvironment. Pancreatic cancer-cell sialylation can promote tumor-associated macrophage differentiation through Siglec-7 and Siglec-9, providing a mechanistic precedent for such interactions, although this mechanism was not tested here^15,43^. Our analysis extends this approach to transcriptional programs initially defined within glycan-selected tumor regions. Given the established prognostic utility of noninvasive serum CA19-9 measurements, deeper molecular characterization of CA19-9-associated tumors and their relationships with clinical outcomes and drug vulnerabilities may provide a practical framework for therapeutic stratification in PDAC^44^.

The drug-response analyses further support the potential translational relevance of the CA19-9-associated programs. Previous studies have linked PDAC transcriptional features to chemotherapy sensitivity in patient-derived organoids and shown that microenvironment-dependent changes in cancer-cell state can alter drug response. In DepMap PDAC cell lines, standard-of-care and other clinically relevant agents exhibited divergent but generally modest associations with program activity, and Cluster 1 did not show consistent sensitivity to conventional agents. Thus, the poor prognosis associated with Cluster 1 dominance in TCGA-PAAD was not accompanied by an obvious therapeutic sensitivity to standard single-agent treatments in monoculture models. This is notable because Cluster_1 dominance was associated with poor survival in TCGA-PAAD, suggesting that the worse clinical outcome of Cluster_1-like tumors could be explained by drug-resistance profile in monoculture drug screens. Thus, the poor survival associated with Cluster 1 dominance may reflect not only aggressive tumor biology, but also the absence of strong, standard-of-care drug sensitvity detectable in current cell-line pharmacologic screens. Broader pharmacologic screening identified stronger program-specific associations and nominated moexipril as a candidate vulnerability in Cluster 1-like cells. This finding is notable given retrospective studies associating renin–angiotensin system inhibitor use with improved outcomes in patients with PDAC^45–48^. Preclinical studies have shown that losartan can reduce stromal barriers and improve chemotherapy delivery, and a single-arm phase 2 trial evaluated losartan with FOLFIRINOX and chemoradiotherapy in locally advanced PDAC^49,50^. These studies provide context for investigating angiotensin-pathway interventions but do not establish the mechanism underlying the moexipril association observed in our monoculture analysis.Although experimental validation is required, the convergence of a Cluster 1-specific pharmacologic association, poor patient survival, and retrospective clinical observations illustrates how glycan-associated transcriptional programs may generate therapeutic hypotheses for clinically adverse PDAC states.

Several limitations should be considered. First, although the GeoMx-derived programs were recovered across independent datasets, the small discovery cohort may not capture the full diversity of transcriptional states associated with CA19-9-positive PDAC. Second, the external datasets lacked direct CA19-9 glycan measurements. Recovery of these programs therefore supports their transcriptional generalizability but does not establish CA19-9 positivity in the corresponding tumors or cells. Third, bulk RNA-seq and Visium spots contain mixtures of cell types, limiting the extent to which projected program scores can be attributed specifically to malignant cells. Fourth, the cell-line drug-response analyses were limited by sample size and monoculture conditions. These associations identify candidate vulnerabilities but do not establish causal dependencies or predict treatment response in patients. Larger spatial cohorts with matched glycan measurements will help define the breadth and glycan specificity of these programs, while functional studies will be needed to validate their associated therapeutic vulnerabilities.

Together, these findings support a model in which CA19-9-positive PDAC contains multiple cancer-cell transcriptional states with distinct clinical and therapeutic associations. By integrating glycan-defined spatial profiling with cross-platform transcriptomic projection, this study provides a framework for resolving heterogeneity within a clinically important glycan-defined PDAC population. We therefore recommend that future PDAC profiling studies integrate multiplex glycan imaging, spatial transcriptomics, clinical outcome annotation, and experimental drug testing to refine these programs and determine whether they can better target cancer-intrinsic mechanisms for clinical benefit.

## Methods

### GeoMx Spatial Transcriptomics

#### Tissue Processing

The tissue samples were collected from surgical resections of patients with PDAC at Duke University Medical Center. All samples were formalin-fixed and paraffin-embedded. The tissue sections used for this study were from tissue blocks not needed for patient evaluation.

Spatial transcriptomics was performed using the nanoString GeoMx® Digital Spatial Profiler. Sections were cut at 6 μm thickness and mounted on plus-charged slides (Epredia Colormark Plus CM-4951WPLUS-001). Slides were baked at 60 °C for 1 hour and stored at 4 °C in a vacuum sealed container containing desiccant for up to two weeks. All subsequent steps were performed using RNase-free conditions and DEPC treated water. Slides were de-paraffinized with 3 sequential 5-minute washes in xylenes, followed by 2 washes in 100% ethanol for 5 minutes, 1 wash in 95% ethanol, and 1 wash in 1x PBS. Target retrieval was performed in target retrieval reagent (10x Invitrogen 00-4956-58 EDTA pH 9.0) diluted to 1x in the BioGenex EZ-Retriever System for 10 minutes at 95 °C. Slides were then washed with 1x PBS for 5 minutes. Slides were then incubated in 0.1 μg/mL proteinase K (Invitrogen 25530-049) for 10 minutes at 37 °C and washed in 1x PBS for 5 minutes at room temperature. Slides were then incubated with hybridization probes (nanoString Cat# 121401103) diluted in Buffer R (provided in the GeoMx RNA Slide Prep FFPE-PCLN kit, catalog # 121300313) and DEPC treated water in a hybridization oven at 37 °C for 16-20 hours.

Following probe incubation, slides were washed with stringent washes (equal parts formamide and 4x SSC buffer) at 37 °C twice for 25 minutes each. Then slides were washed twice in 2x SSC buffer. Slides were incubated in 200 μL buffer W (provided in the GeoMx RNA Slide Prep FFPE-PCLN kit, catalog # 121300313) for 30 minutes and incubated in protein probe (GeoMx 10 Proteome Atlas 500 (IPA)Kit, #531-121300160) and morphology markers mixed in Buffer W ( GATA3 Antibody (HG3-31): sc-268AF647 diluted 1:100, CYP1A1 Antibody (B-4): sc-25304AF594 diluted 1:150, Pan Cytokeratin Monoclonal Antibody (AE1/AE3), Alexa Fluor™ 488 diluted 1:100) at 4 °C overnight. Slides were washed in 1x TBST three times for 10 minutes each wash followed by post fixation in 4% PFA (Invitrogen FB002) for 30 minutes. Following post fixation, slides were washed in 1x TBST two times for 10 minutes each wash. Slides were then incubated with nuclei marker Syto83 (Thermo Scientific S11364, 1:1000) in Buffer W for 1 hour at room temperature in a humidified chamber. Slides were washed in 1x TBST two times for 10 minutes each and placed in the nanoString GeoMx® DSP instrument. Syto83 immunofluorescence was utilized for autofocus of GeoMx imaging.

#### NGS library prep and sequencing

The tissue areas of interest are located using fluorescent imaging on the GeoMx instrument. Regions of interest (ROIs) are selected based on morphology marker stainings and segmented into areas of interest (AOIs). AOIs are processed sequentially on the instrument using UV light focused through each AOI, releasing the photocleavable oligos into wells of a 96-well plate. Of note, the first plate of AOIs was collected on day 3, but due to instrument failure, the slide was stored overnight, and the second plate of AOIs were re-drawn and collected on day 4. After AOI collection, indexing primers are hybridized during Nanostring library preparation. Quality and quantity of the finished library pools were assessed using a combination of Agilent DNA High Sensitivity chip (Agilent Technologies, Inc.) and QuantiFluor® dsDNA System (Promega Corp.). Paired end, 27bp sequencing will be performed on an Element AVITI sequencer (Element Biosciences, San Diego, CA, USA). Base calling is performed on instrument and AVITI OS (v2.6.2) output are demultiplexed and converted to FastQ format with Element Biosciences Bases2fastq v1.7.0. Fastq files were then converted to Digital count conversion files (DCC) using GeoMx NGS Pipeline v3.1.1.6.

#### Quality Control

Digital count-conversion files and sample annotations were imported into R as NanoStringGeoMxSet objects using NanoStringNCTools and GeoMxTools. RNA and protein data were processed separately, and no-template control segments were removed before quality assessment.

Areas of illumination (AOIs) were evaluated using sequencing, alignment, morphology, and probe-level metrics. The prespecified segment-level criteria were a minimum of 1,000 sequencing reads, ≥80% trimmed reads, ≥80% stitched reads, ≥75% aligned reads, and ≥50% sequencing saturation. AOIs were additionally required to have a minimum area of 50 µm². No minimum nuclear-count threshold was imposed. Negative-control counts were required to be ≥1, and the maximum no-template-control count was 9,000.

RNA probes were evaluated using a minimum probe count of 10, a minimum probe ratio of 0.1, and a Grubbs outlier test with α=0.01. Probe sets for which more than 20% of probes failed the outlier test were flagged for exclusion. The limit of quantification (LOQ) was calculated from negative-probe measurements using an LOQ multiplier of 1. Genes were retained when their expression exceeded the LOQ in at least 8% of AOIs, and AOIs were retained when at least 3% of measured features exceeded the LOQ. A high-count cutoff of 10,000 was used for QC reporting. The synthetic negative feature NegProbe-WTX was removed before downstream expression analyses.

Protein quality was assessed using the manufacturer-designated housekeeping proteins and IgG negative controls. For each AOI, background was estimated as the geometric mean of the IgG-control counts after addition of a pseudocount; the arithmetic mean was used if the geometric-mean estimate was nonfinite or nonpositive. Protein signal-to-background ratios were calculated by dividing each protein count by the corresponding AOI-specific background and were inspected on a log2 scale. Raw, normalized, and log-transformed protein-count distributions were also examined across AOIs. Protein AOIs were restricted to those that passed the RNA QC procedure, ensuring matched RNA and protein profiles for integrated analyses.

#### Data and Statistical Analysis

A total of 172 AOIs from three tissue specimens were available before quality control: 82 from BRPC_17-543, 43 from BRPC_19-296, and 47 from BRPC_20-281. Following application of the prespecified RNA quality-control criteria, 164 AOIs were retained, including 75 from BRPC_17-543, 43 from BRPC_19-296, and 46 from BRPC_20-281. Therefore, 7 AOIs were excluded from BRPC_17-543 and 1 AOI was excluded from BRPC_20-281. The protein dataset was restricted to the same AOIs retained after RNA quality control, resulting in matched RNA and protein measurements for all 164 AOIs.

GeoMx RNA counts passing quality control were upper-quartile normalized and log2 transformed for downstream analyses. Protein counts below 1 × 10^−10 were set to 1 and normalized to the 75th percentile using GeoMxTools; normalized protein values were subsequently log2 transformed. Principal component analysis, unsupervised hierarchical clustering, and heatmaps were generated from the normalized expression matrices. Four RNA expression groups were defined by cutting the AOI dendrogram into four clusters. These clusters represent expression programs observed in the discovery specimens and were therefore interpreted as patient-derived CA19-9-associated programs rather than universal PDAC subtypes.

Differential expression among RNA clusters was evaluated using gene-wise linear mixed-effects models implemented with GeoMxTools. Cluster identity was included as a fixed effect. Models containing either AOI or tissue/patient identity as a random intercept were examined, with the tissue-level random-effect model used to evaluate expression differences after accounting for repeated measurements and between-patient variation.

Pairwise contrasts were obtained from estimated marginal means. P values were adjusted separately within each cluster comparison using the Benjamini–Hochberg false-discovery-rate (FDR) procedure. Effect estimates were oriented so that positive values represented enrichment in the indicated cluster. Cluster-associated marker sets were derived by summarizing each gene’s oriented effect across comparisons between the focal cluster and the other clusters and retaining genes meeting the prespecified effect-size and FDR criteria. Gene Ontology enrichment analysis was subsequently performed on the cluster-associated gene sets.

To visualize expression patterns after removal of patient-level effects, a separate linear mixed-effects model containing tissue identity as a random intercept was fit to each gene, and the model residuals were used for patient-adjusted heatmaps and dimension-reduction analyses. Published Classical and Basal-like/Squamous PDAC signatures and glycosylation-associated gene sets were evaluated by extracting the corresponding genes present in each expression dataset and summarizing or visualizing their normalized expression. These signature analyses were descriptive unless otherwise specified.

For validation in TCGA-PAAD, upper-quartile-normalized FPKM values were obtained through TCGAbiolinks. For each tumor and each GeoMx-derived cluster, a program score was calculated as the mean expression of the available marker genes assigned to that cluster. Overall profile identity was summarized using both the mean of the four program scores and the minimum of the four scores. Tumors with both values at or above their respective 75th percentiles were classified as high-profile-identity tumors. Within high-profile-identity tumors, the four program scores were scaled within samples and hierarchically clustered to identify the relatively dominant expression program.

Overall survival was defined as days to death for deceased patients and days to last follow-up for living patients. When multiple aliquots were available for a patient, the sample with the highest mean program score was retained, yielding one observation per patient. Survival distributions were estimated using the Kaplan– Meier method and compared using two-sided log-rank tests. Cox proportional-hazards models were used to estimate associations between program-defined groups and survival.

Independent PDAC Visium data were processed in Seurat. Cancer-enriched spots were identified by comparing cancer, noncancer, and immune module scores derived from published cell-type gene sets, followed by spatial smoothing of the cancer score and threshold-based classification. GeoMx program scores were calculated for each cancer-enriched spot as the mean normalized expression of the available marker genes in each cluster. A spot was classified as profile-high when its mean score across the four programs exceeded 2.214, corresponding to the prespecified third-quartile cutoff used in the analysis. Samples in which at least 50% of evaluated spots were profile-high were classified as high-fraction samples. Because the four absolute program scores were strongly correlated, relative program scores and the program with the largest relative score were used to characterize dominant program emphasis. Correlations among program scores were calculated using Pearson correlation with pairwise-complete observations.

For cell-line analyses, raw RNA-sequencing counts were aggregated by gene symbol, normalized for library size with DESeq2 size factors, and variance-stabilized using a blind variance-stabilizing transformation for visualization and clustering. For DepMap analyses, pancreatic cancer cell lines with both transcriptomic and pharmacologic-response data were retained. For each cell line, GeoMx-derived program scores were calculated as the mean expression of the available marker genes. Relative program scores were calculated by subtracting the cell line’s mean score across all four programs from each individual program score.

Drug-response values were modeled as a function of each continuous relative program score using separate ordinary least-squares regressions. Only drug–program combinations represented by at least six distinct cell lines were analyzed. Regression slopes, nominal P values, and coefficients of determination (R²) were reported. P values were adjusted within each program across tested drugs using the Benjamini–Hochberg procedure. In the DepMap response metric, more negative values represent greater drug sensitivity; consequently, a negative regression coefficient indicates that increasing program activity is associated with greater sensitivity, whereas a positive coefficient indicates reduced sensitivity or relative resistance.

Categorical comparisons based on dominant program identity were assessed using Kruskal–Wallis tests, with pairwise Wilcoxon rank-sum tests used for exploratory follow-up comparisons when groups contained at least two observations.

Power for continuous drug-response models was evaluated using the noncentral F distribution for a single-predictor linear model. The number of cell lines required for 80% power was estimated under both a nominal two-sided α of 0.05 and a Bonferroni-adjusted α based on the number of modeled associations. Given the limited number of cell lines and the number of drug–program comparisons, these pharmacogenomic analyses were considered exploratory; interpretation emphasized effect size, direction, R², consistency across programs, and FDR-adjusted evidence rather than nominal significance alone. Unless otherwise stated, tests were two-sided, nominal statistical significance was defined as P<0.05, and multiplicity-adjusted findings were evaluated at FDR<0.05. Analyses were performed in R using packages including NanoStringNCTools, GeoMxTools, Biobase, lme4, DESeq2, TCGAbiolinks, Seurat, survival, survminer, and associated visualization packages.

### Multiplex Immunofluorescence (mIF) Staining

Multiplex immunofluorescence staining was performed on the same tissue sections used for the spatial transcriptomics. For each staining cycle, sections were incubated with fluorophore-conjugated primary antibodies overnight at 4°C, followed by counterstaining with DAPI for 5 minutes to visualize nuclei. Images were acquired after each cycle using an Axio Scan 7 automated slide scanner (Carl Zeiss Microscopy, Oberkochen, Germany). Before the subsequent cycle of staining, the existing fluorescence signal was extinguished by treating the sections with 6% hydrogen peroxide in a 250 mM carbonate buffer (pH 9.5) for 20 minutes. The following primary antibodies were utilized for the mIF panel: CA19-9 (clone 1116-NS-19-9, MyBioSource), TRA-1-60 (clone TRA-1-60, Novus Biologicals), and anti-Tn antigen (clone reBaGs6, Beth Israel Deaconess Medical Center).

### Human Pancreatic Tissue

The study was conducted under protocols approved by the Institutional Review Boards at the Van Andel Research Institute, and the Duke University Medical Center. All subjects provided written, informed consent, and all methods were performed in accordance with the relevant guidelines and regulations. Human pancreatic cancer tissue samples were obtained from surplus surgical resections. Tissues were immediately fixed in 10% neutral buffered formalin (NBF) at room temperature for 24 h. Following fixation, tissues were trimmed to a thickness of 3–4 mm, placed into embedding cassettes, and processed overnight through a graded ethanol series, cleared in xylene, and infiltrated with paraffin wax using an automated tissue processor. The specimens were then embedded into paraffin blocks, chilled on ice, and sectioned at a thickness of 6 µm using a rotary microtome. Finally, sections were floated on a 40–42°C water bath, mounted onto positively charged glass slides, and baked at 60°C for 1–2 h to ensure optimal tissue adherence prior to downstream histological analysis.

## Data and Code Availability

Data and code will be made publicly available upon publication.

## Acknowledgements

The author(s) thank the Van Andel Spatial Transcriptomics Core ([RRID:SCR_027861](https://coremarketplace.org/?FacilityID=5718)) and Pathology and Biorepository Core (RRID:SCR_022912) for providing tissue processing, spatial transcriptomics and sequencing facilities and services. Some figures were made using Biorender.

---

**Supplemental Figure 1.**
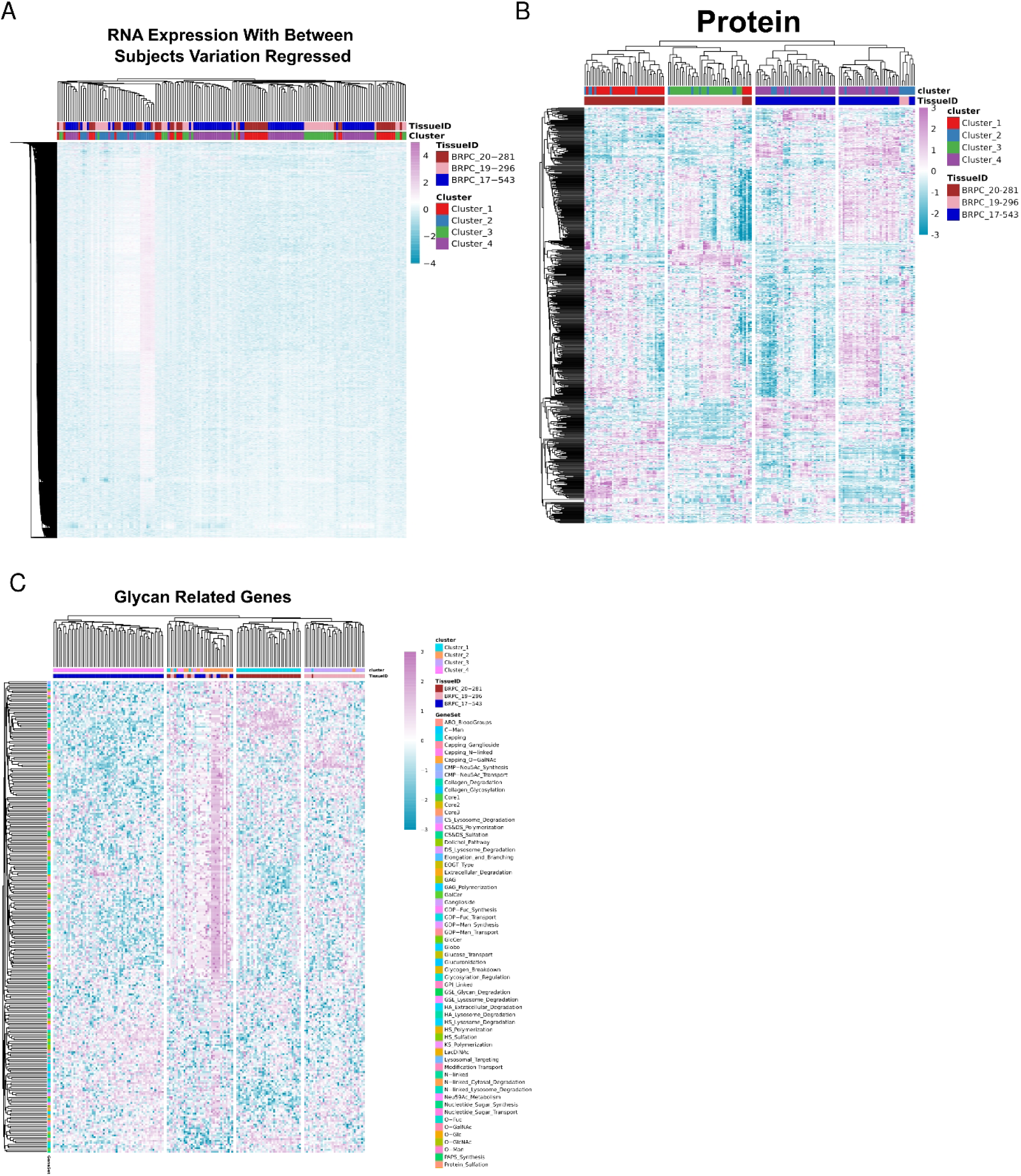
Patient-adjusted RNA, protein, and glycosylation-related expression patterns in CA19-9–positive PDAC. **(A)** Unsupervised hierarchical clustering of GeoMx RNA expression after accounting for between-patient variation. A gene-wise linear mixed-effects model containing tissue identity as a random intercept was used to remove patient-level effects, and the resulting residual expression values are shown. **(B)** Unsupervised hierarchical clustering of normalized GeoMx protein expression across matched areas of illumination (AOIs), demonstrating structured protein-level variation among the RNA-defined expression programs. (C) Unsupervised hierarchical clustering of normalized expression for glycosylation-associated genes, showing coordinated differences in glycan-related transcriptional pathways across the same sample and program architecture. In all panels, columns represent GeoMx AOIs and rows represent measured genes or proteins. Column annotations indicate tissue identity and RNA-cluster assignment; the row annotation indicates the associated glycosylation pathway or functional gene set. Colors represent relative expression according to the scales shown.

**Supplemental Figure 2.**
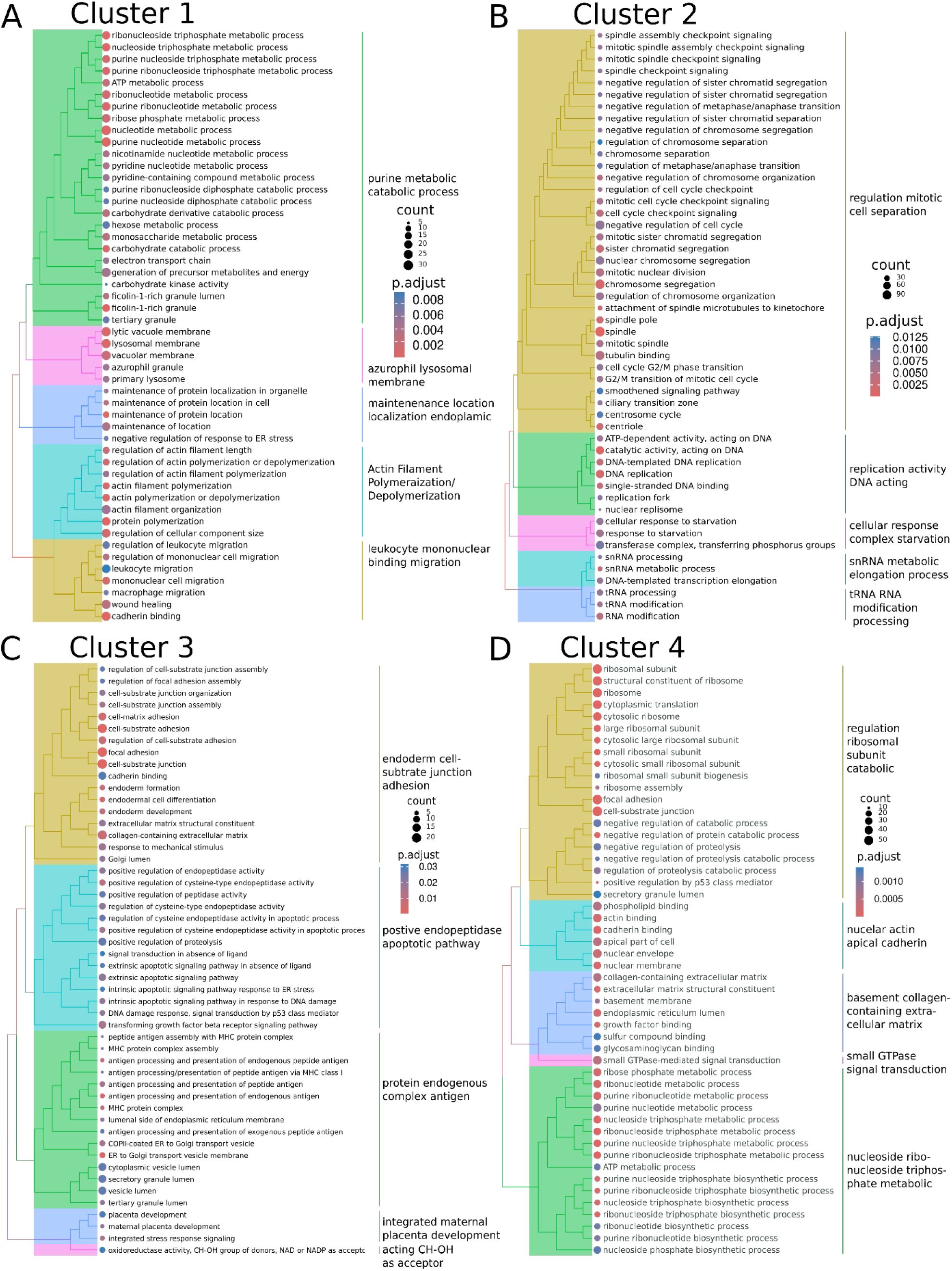
Gene Ontology analysis of the GeoMx-derived CA19-9-associated transcriptional programs. Tree plots show enriched Gene Ontology Biological Process terms associated with marker genes for **(A)** Cluster 1, **(B)** Cluster 2, **(C)** Cluster 3, and **(D)** Cluster 4. Enrichment was performed using clusterProfiler with Benjamini–Hochberg false-discovery-rate correction; terms with *q*<0.05 were retained. Up to the 50 highest-ranking terms are displayed and grouped by pairwise semantic similarity using average-linkage clustering. Node size represents the number of cluster-marker genes assigned to each term, node color represents the adjusted *P* value, and branch colors denote groups of semantically related terms. Cluster 1 was enriched for inflammatory, leukocyte-migration, wound-healing, and cytoskeletal-remodeling processes; Cluster 2 for DNA replication, G2/M progression, nuclear division, and chromosome segregation; Cluster 3 for antigen processing and presentation, differentiation, TGF-β signaling, and stress-response processes; and Cluster 4 for ribosomal assembly and cytoplasmic translation.

**Supplementary Figure 3.**
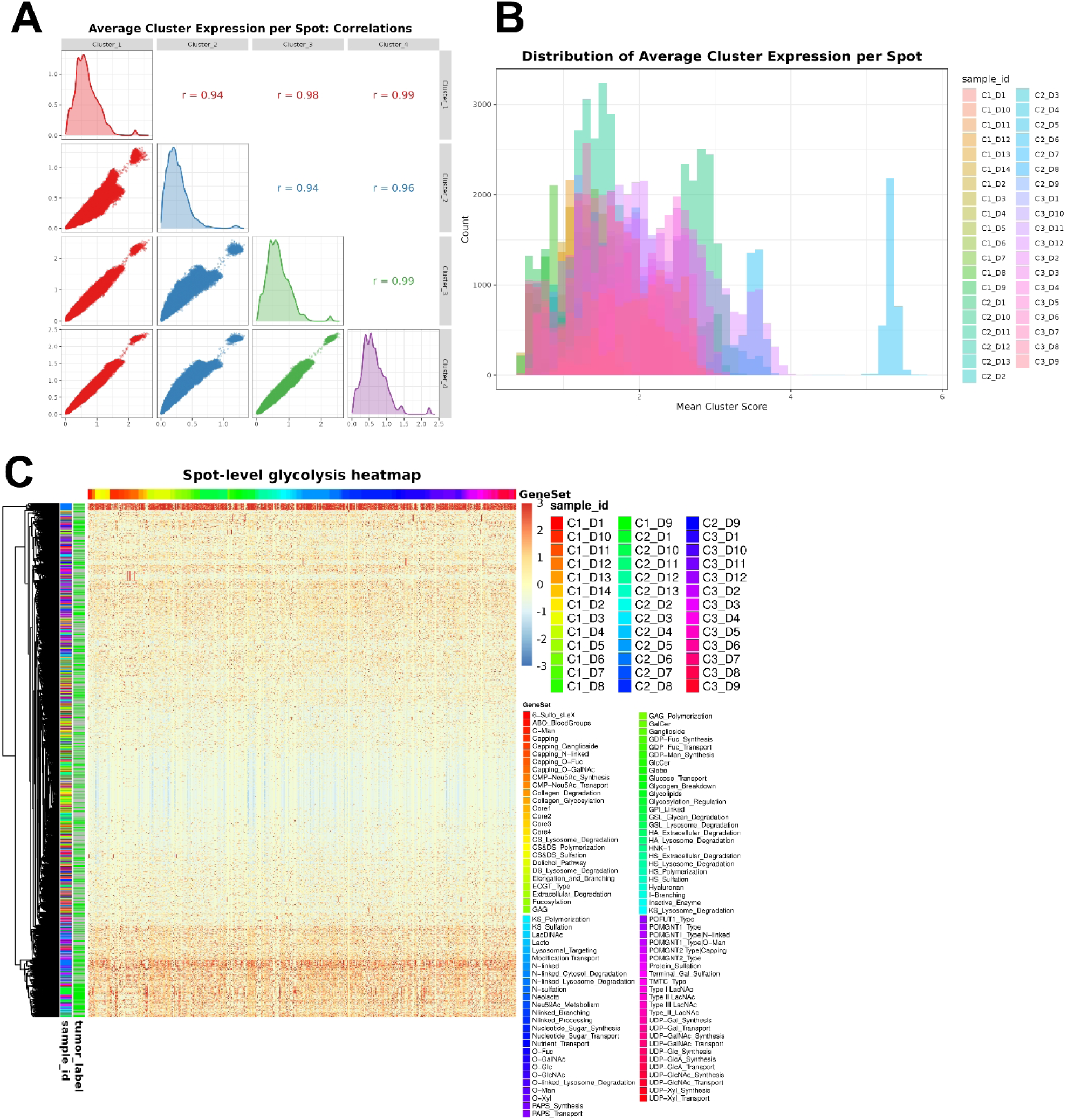
GeoMx-derived program scores and glycosylation-associated gene expression in the independent PDAC Visium cohort. **(A)** Pairwise relationships among the four GeoMx-derived program scores across Visium spots. Each program score was calculated as the mean normalized expression of its available marker genes. Diagonal plots show score distributions, lower-triangle plots show spot-level pairwise relationships, and upper-triangle values report Pearson correlation coefficients calculated using pairwise-complete observations. Scores were strongly positively correlated across programs (r = 0.94– 0.99). **(B)** Overlaid histograms showing the distribution of the mean program score per spot, colored by Visium sample. The mean score summarizes expression across the four GeoMx-derived programs; a threshold of >2.214 was used to designate high-scoring spots for the sample-level analysis in Figure 3D. **(C)** Heatmap of glycosylation-associated gene expression across randomly sampled Visium spots (up to 200 per sample). Rows represent spots ordered by hierarchical clustering, and columns represent genes grouped by annotated gene set. Side annotations indicate sample identity and tumor classification; the upper annotation indicates gene-set membership.

**Supplemental Figure 4.**
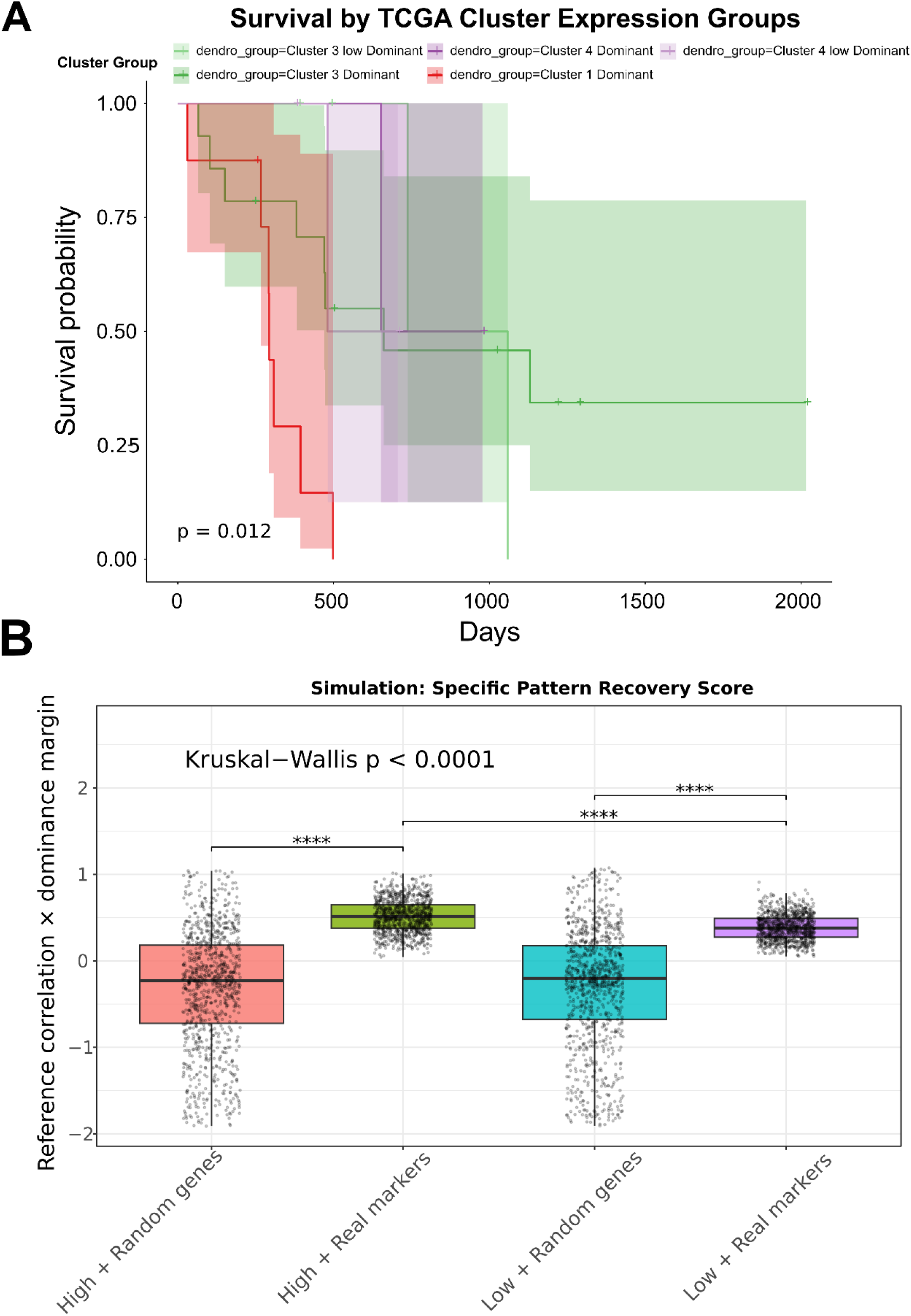
Survival and simulation-based evaluation of GeoMx-derived program structure in TCGA-PAAD. **(A)** Kaplan–Meier overall-survival curves for high-profile-identity TCGA tumors stratified by the dominant-program groups obtained from hierarchical clustering of within-sample-scaled GeoMx program scores. Survival time was defined as days to death or last follow-up, with one sample retained per patient. Shaded regions indicate 95% confidence intervals, censoring is indicated by tick marks, and the displayed *P* value was calculated using a two-sided log-rank test. **(B)** Simulation-based assessment of the specificity and reproducibility of the GeoMx-derived marker structure. In each of 1,000 iterations, three tumors were randomly selected from either the high-profile-identity or remaining TCGA sample pool and scored using either the authentic GeoMx marker sets or size-matched random gene sets that excluded the authentic marker genes. Program scores were scaled within each tumor. Pattern-recovery scores were calculated as the product of the mean correlation with the high-profile-identity reference pattern and the mean dominance margin between the highest and second-highest program scores. Points represent individual simulations. Differences among the four conditions were assessed by a Kruskal–Wallis test, followed by pairwise Wilcoxon rank-sum tests with Benjamini–Hochberg correction for the indicated comparisons (All comparisons p<0.0001).

**Supplemental Figure 5.**
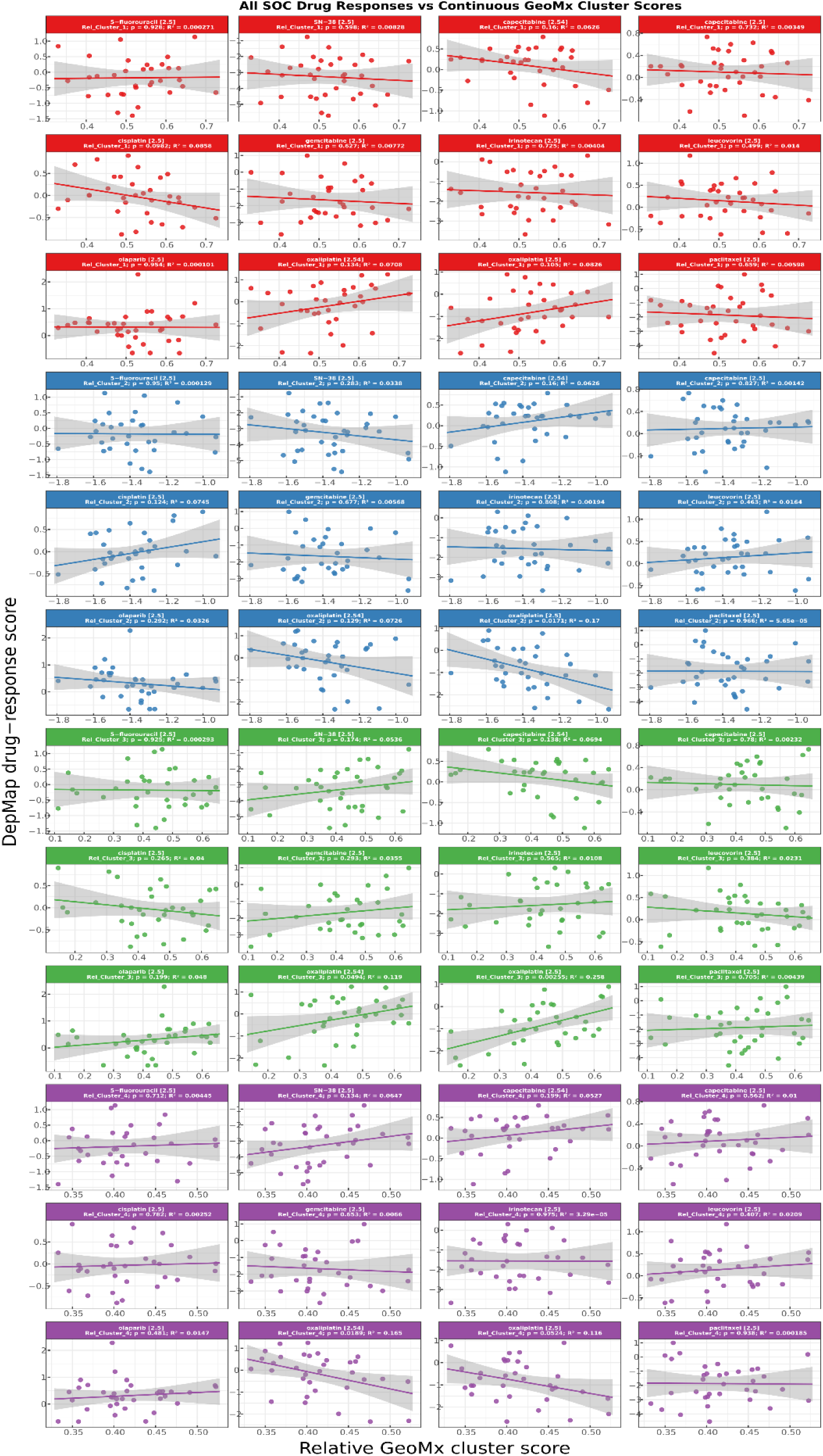
Standard of Care Cluster Program Drug Response Associations. Associations between GeoMx-derived transcriptional programs and in-vitro response to PDAC standard-of-care components and related agents. Scatterplots show associations between relative Cluster 1, Cluster 2, Cluster 3, and Cluster 4 program scores and individual drug–dose measurements selected from the DepMap PRISM screen. Agents were selected when their drug name, compound annotation, mechanism of action, or molecular target matched a component of established PDAC treatment or a clinically relevant related agent, including fluorouracil, gemcitabine, irinotecan or its active metabolite SN-38, paclitaxel, oxaliplatin, capecitabine, leucovorin, olaparib, and cisplatin. Each point represents a pancreatic cancer cell line. Lines and shaded regions show ordinary least-squares regression estimates and 95% confidence intervals, respectively. Facet labels identify the drug and experimental concentration and report the modeled program, nominal *P* value, and coefficient of determination (R²); facet colors denote the corresponding GeoMx program. The response variable is the PRISM replicate-collapsed log-fold change in cell viability relative to DMSO. More negative values indicate greater in-vitro sensitivity; therefore, negative slopes indicate increasing sensitivity with increasing program score, whereas positive slopes indicate reduced sensitivity or relative resistance. Models were limited to drug–program combinations represented by at least six distinct cell lines. Benjamini–Hochberg false-discovery-rate correction was calculated across drug–dose tests within each program, although the facet labels display nominal *P* values. These exploratory single-agent, single-concentration associations do not model multidrug regimens, clinical dosing, or patient treatment response.

**Supplemental Figure 6.**
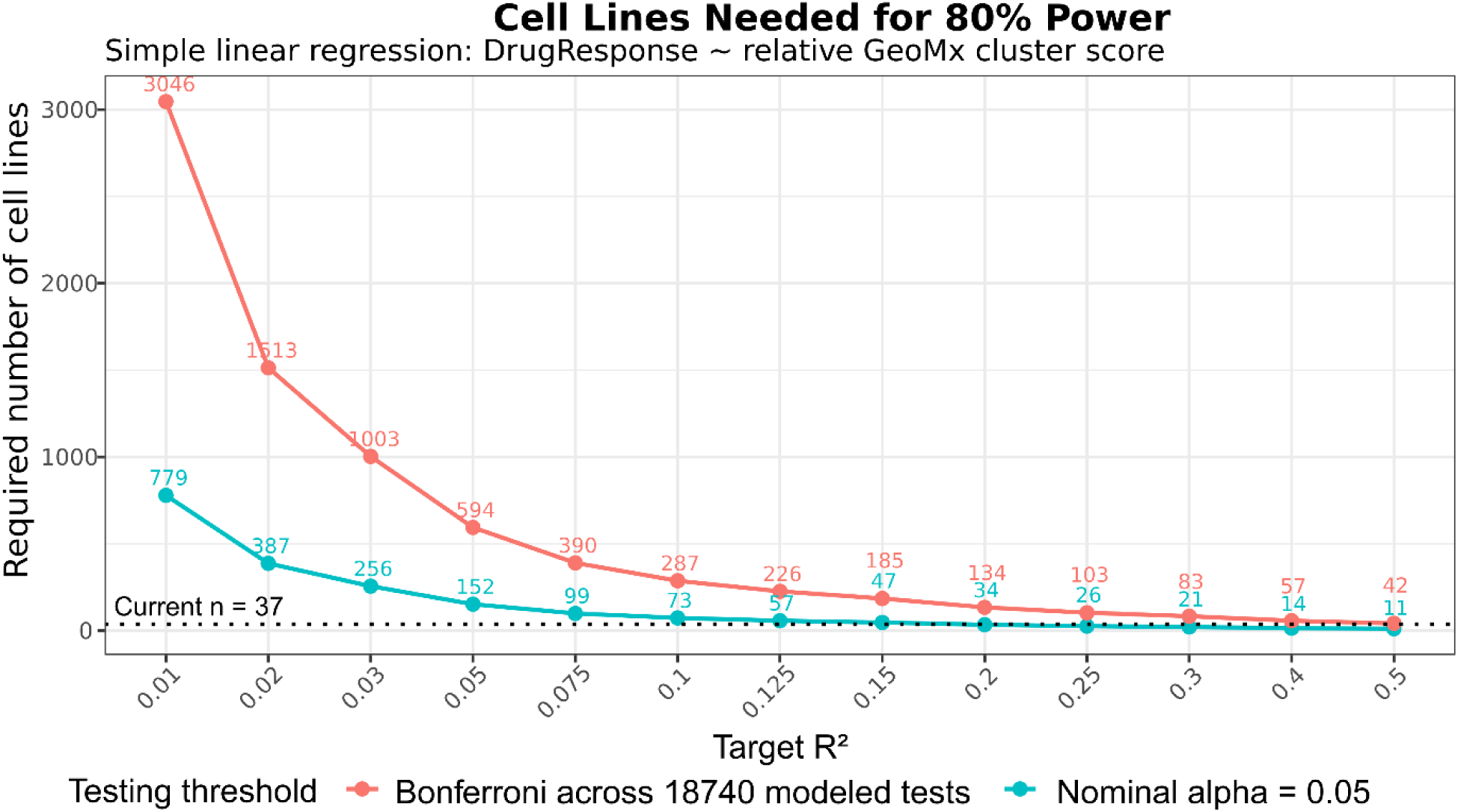
Power Calculation For Drug Response Association Calculations. Estimated number of cell lines required to achieve 80% power for single-predictor linear regression across a range of target effect sizes expressed as R². Curves compare a nominal two-sided α=0.05 with a Bonferroni-adjusted α based on the number of modeled drug–program associations. The horizontal dotted line indicates the 37 cell lines available in the matched dataset; numbers denote the estimated sample size required at each target R².

